# Artificial lighting affects the predation performance of the predatory bug *Orius insidiosus* (Say) against the Western flower thrips *Frankliniella occidentalis* (Pergande)

**DOI:** 10.1101/2024.10.01.616144

**Authors:** Morgane L. Canovas, Paul K. Abram, Jean-François Cormier, Tigran Galstian, Martine Dorais

## Abstract

Protected crops, such as greenhouses and indoor farming environments, using light-emitting diodes (LEDs) enable the modulation of the light spectrum, intensity, and photoperiod for agronomic purposes. This creates dynamic artificial light conditions for insects and arachnids, including predators used in biological control. Despite increasing interest, the effects of LEDs on predator behavior and control performance remain poorly understood. In the laboratory, we examined the locomotion and predation behaviors of the generalist predator *Orius insidiosus* against the pest *Frankliniella occidentalis* under various light spectra and intensities. We tested narrowband blue, green, and red spectra, three ratios of red and blue light, and a spectrum combining all three colors across a gradient of light intensity in microcosms. *Orius insidiosus* was active and successfully attacked prey under all lighting conditions, with 70% of individuals engaging in predation during the observation period. The light spectrum significantly influenced all recorded behaviors, while light intensity had negligible effects. Narrowband spectra led to the highest attack probabilities, but the mixed blue-red spectrum with a higher proportion of red light yielded the highest prey capture rates. The spectrum with all three colors showed intermediate capture success. These trends were consistent regarding capture probability in more complex environments with cucumber plants, where thrips were exposed to 24-hour artificial light sequences before and during predator releases. However, thrips survival rates remained similar across all lighting treatments. Our results demonstrated that while lighting treatments affect predator behavior, *Orius insidiosus* retains its ability to capture prey under various light conditions. This paves the way for developing lighting strategies that balance plant productivity with effective biological control in protected crop environments.

## Introduction

Artificial lighting is critical for crop production in protected environments (i.e., greenhouses, tunnels, indoor farming) because it supports plant productivity when solar radiation is insufficient in the northern latitudes (Dorais et al. 2017), or represents the sole light source available for fully indoor cultivation (Cary & Stutte 2017). The widespread adoption of light-emitting diodes (LEDs) allows precise manipulation of photoperiod, light intensity, and spectrum, enabling year-round plant production. However, sensitivity to different parts of the electromagnetic spectrum varies between plants and the pest and beneficial insects that inhabit protected plant production environments. The insect community in protected crops includes arthropods used for augmentative biological pest control (Van Lenteren et al. 2018). In general, plants preferentially absorb wavelengths in the blue (400–500 nm) and red (600–700 nm) ranges for photosynthesis, as these correspond to the absorption peaks of chlorophyll *a* (430 and 663 nm) and *b* (453 and 642 nm) (Bantis et al. 2018). Consequently, artificial LED lighting that contains both red and blue wavelengths is commonly used in horticulture. On the other hand, insects generally exhibit dichromatic sensitivity to light, with absorption peaks in the UV-blue (λ_max_∼350 nm) and green (λ_max_ ∼530 nm) ranges (Briscoe & Chittka 2001; Johansen et al. 2011). Compound eyes and ocelli enable insects to perceive variations in duration of illumination, light intensity, spectra, and polarization (Gullan & Cranston 2014). The integration of this information drives their behaviors and life history traits, such as resource location and development (Johansen et al. 2011). Manipulating light environments thus has the potential to affect the efficacy of pest control in protected crops by affecting both on pests and their natural enemies. However, the application of this knowledge in horticultural contexts has predominantly targeted pests (e.g., Shimoda & Honda 2013); for example, to develop trapping strategies (Ben-Yakir 2020) or manipulate insect behavior (Tyler-Julian et al. 2018; Ben-Yakir 2020).

Until quite recently, there was very little known about the visual sensitivity of arthropods used in biological control to the different light spectra applied in protected crops (Vänninen et al. 2010; Johansen et al. 2011). The use of visual cues to enhance beneficial insect activity has mainly focused on stimulating their recruitment from natural habitats into greenhouses or fields (Shimoda & Ben-Yakir 2020). However, recent studies have shown that commercially available parasitoid species do, in fact, respond to photoperiod and spectral quality, particularly different ratios of blue and red light. For *Aphidius ervi* (Haliday), an unfavorable light environment can induce a male-biased sex ratio (Cochard et al. 2019) or trigger diapause (Tougeron et al. 2019), potentially compromising pest control effectiveness. On the other hand, for the aphid parasitoid *Aphidius matricariae* (Haliday), total locomotor activity appears similar across artificial lighting treatments (Fraser et al. 2023a), and its efficacy as a biological control agent against *Myzus persicae* (Sulzer) was equivalent across different lighting treatments involving supplementation with broad-spectrum white or narrowband red and blue spectra (Fraser et al. 2023b). The behavioral responses of the many other arthropod parasitoids and predators introduced in augmentative biological control to various artificial light spectra used in protected agriculture remain unknown.

Heteropteran predators are an important group of natural enemies used for biological control in protected crops (Van Lenteren 2012). This includes bugs in the *Orius* genus (Hemiptera: Anthocoridae), which are sold commercially worldwide (Van Lenteren 2012; Van Lenteren et al. 2018). *Orius insidiosus* (Say) is widely used in Europe, North America, and South America, where it has been used for biological control of plant pests for around forty years (Van Lenteren 2012). It is particularly known for its efficiency in controlling thrips, including the cosmopolitan pest *Frankliniella occidentalis* (Pergande) (Reitz et al. 2020). *Orius insidiosus* is introduced in both vegetable and ornamental crops and can thus be exposed to a wide range of lighting regimes optimized for horticulture, including various red-blue light ratios and broader spectra incorporating blue, green, and red wavelengths. While the response of its primary target host, *F. occidentalis*, to lighting is relatively well understood across different contexts (Lopez-Reyes et al. 2022; Dearden et al. 2024; Grupe & Meyhöfer 2024), previous studies on the photobiology of *O. insidiosus* have mainly focused on manipulating photoperiod to address the issue of winter diapause in protected crops (Kingsley & Harrington 1982; Van Den Meiracker 1994; Stack & Drummond 1995; 1997; Ruberson, Shen & Kring 2000; Herrick et al. 2021). However, *Orius* species’ prey orientation, dispersal, locomotor activity, and reproduction also appear to be sensitive to both light spectrum and intensity (Henaut et al. 1999; Legarrea et al. 2012; 2012; Wang et al. 2013; Ogino et al. 2015; 2016; 2020; Park et al. 2023). Despite the identification of positive effects of artificial lighting on diapause avoidance and the attraction of *Orius* spp., the potential for horticultural lighting to enhance its predation efficiency for biological control in protected crops has not been investigated. Additionally, studies on the behavior of *Orius* spp. in response to light spectra and intensity have traditionally been conducted separately, even though these two parameters vary together in nature and can be simultaneously controlled using LEDs. Finally, no specific information is available regarding the predation capabilities of *Orius insidiosus* under different light spectra and intensity conditions, although this species is widely used as a natural enemy under artificial lighting at northern latitudes.

In this study, we tested the effects of the light spectrum and intensity on the small-scale predation behaviors of *Orius insidiosus* towards its prey, the pest thrips *Frankliniella occidentalis*. A custom optical device allowed us to assess the predator’s ability to capture prey under a wide array of different lighting environments. We then examined whether these trends persisted in a more complex environment (including plants) using a subset of the lighting treatments. Given insects’ dichromatic sensitivity peaks in the blue and green spectra (Briscoe & Chittka 2001), we predicted that these spectra would result in the highest predation performance. Conversely, based on previous studies of another *Orius* species (Wang et al. 2013), we anticipated that exposure to red light would induce stress. Specifically, we predicted that red light would trigger an avoidance response, characterized by relatively high locomotor activity and a reduced frequency of successful captures. Our goal was to improve understanding of whether the light spectra commonly used in horticulture are conducive to biological control by *O. insidiosus*, and whether lighting regimes in protected crops need to be optimized to support both predation and plant productivity.

## Materials and methods

### Insect rearing

The pest thrips, *F. occidentalis*, were sourced from a permanent colony maintained in growth chamber at Laval University (CONVIRON, Winnipeg, Manitoba, CA) set at 25 ± 2° C, 65 ± 2% RH, and a photoperiod of 16:8 h light: dark with standard fluorescent lighting (Sylvania F72T12/CW/VHO 160W). Introductions of new *F. occidentalis* individuals originating from university greenhouses or commercial garden centers were conducted multiple times annually. The identity of new individuals was verified under a stereomicroscope (OLYMPUS, Quebec, Quebec, CA; model SZ61) using the identification key by Mound & Kibby (1998) before they were introduced into the colony. Rearing was conducted in transparent 1 L Mason glass jars, with the lid replaced by Nitex (160 µm nylon mesh screen). The jars were filled with around 2 cm of vermiculite, and six commercial green bean pods served as both a food source and oviposition substrate. New pods were introduced weekly after being disinfected for 10 minutes in a 5% dilution of bleach solution (LAVO PRO 6) (Labbé et al. 2018).

*Orius insidiosus* predatory bugs were obtained from a commercial biological control company (©ANATIS Bioprotection, Saint-Jacques-le-Mineur, Quebec, CA) for periodic rearing during the trials. Upon receipt, *O. insidiosus* adults, aged approximately 2 to 3 days, were placed in transparent 1 L Mason jars with the lid replaced by Nitex, at a rate of 500 individuals per jar. Each jar contained approximately 3–4 cm of a mixture of buckwheat husks and vermiculite, as well as a piece of absorbent paper, to create a complex environment and reduce cannibalism among predators while ensuring suitable humidity conditions (Schmidt et al. 1995). Six commercial green bean pods, previously disinfected in the same manner as for thrips rearing, provided a water source and oviposition substrate, while six 2 cm diameter stickers coated with frozen *Ephestia khuniella* eggs (©ANATIS Bioprotection) served as the primary food source (Labbé et al. 2018). Green bean pods and *E. khuniella* egg sources were replaced every 3 days. Predators were maintained in growth chamber (CONVIRON, Winnipeg, Manitoba, CA; 25 ± 2° C, 65 ± 2% RH, and a photoperiod of 16:8 h light: dark with standard fluorescent lighting ©SYLVANIA F72T12/CW/VHO 160W).

### Lighting device and treatments

To investigate the isolated effects of key spectral ranges relevant to the photosensitivity of insects and plants, we tested narrowband wavelength spectra, as well as blue and red ratios typically used in horticulture, along with an ‘extended’ spectrum containing blue, green, and red light (Table 1). The composition of the extended spectrum included these three colors in proportions similar to those previously identified by Garcia & Lopez (2020) to promote growth in greenhouse cucumber transplants. During preliminary trials, exposure to UV light (λ_max_ = 400 nm) resulted in no captures and was thus not considered further (Supplementary material 1). These various spectra were tested in combination with a gradient of light intensity ranging from 50 to 600 µW/cm^2^, resulting in a total of 35 tested spectrum × light intensity combinations (Table 1). The light intensity range was determined by a trade-off between video recording quality and the specifications of our optical device, which are described in the following paragraph. For context, the target artificial light intensity in commercial vegetable greenhouses across the province of Quebec ranges from 80 to 300 µmol/m^2^/s (Sébastien Couture, agr., CLIMAX Conseil, personal communication), while it is 42W in *O. insidiosus* rearing facilities (Dr. Silvia Todorova, ANATIS Bioprotection, personal communication). Our intensity treatments were below these intensity values (Table 1).

**Table 1.**
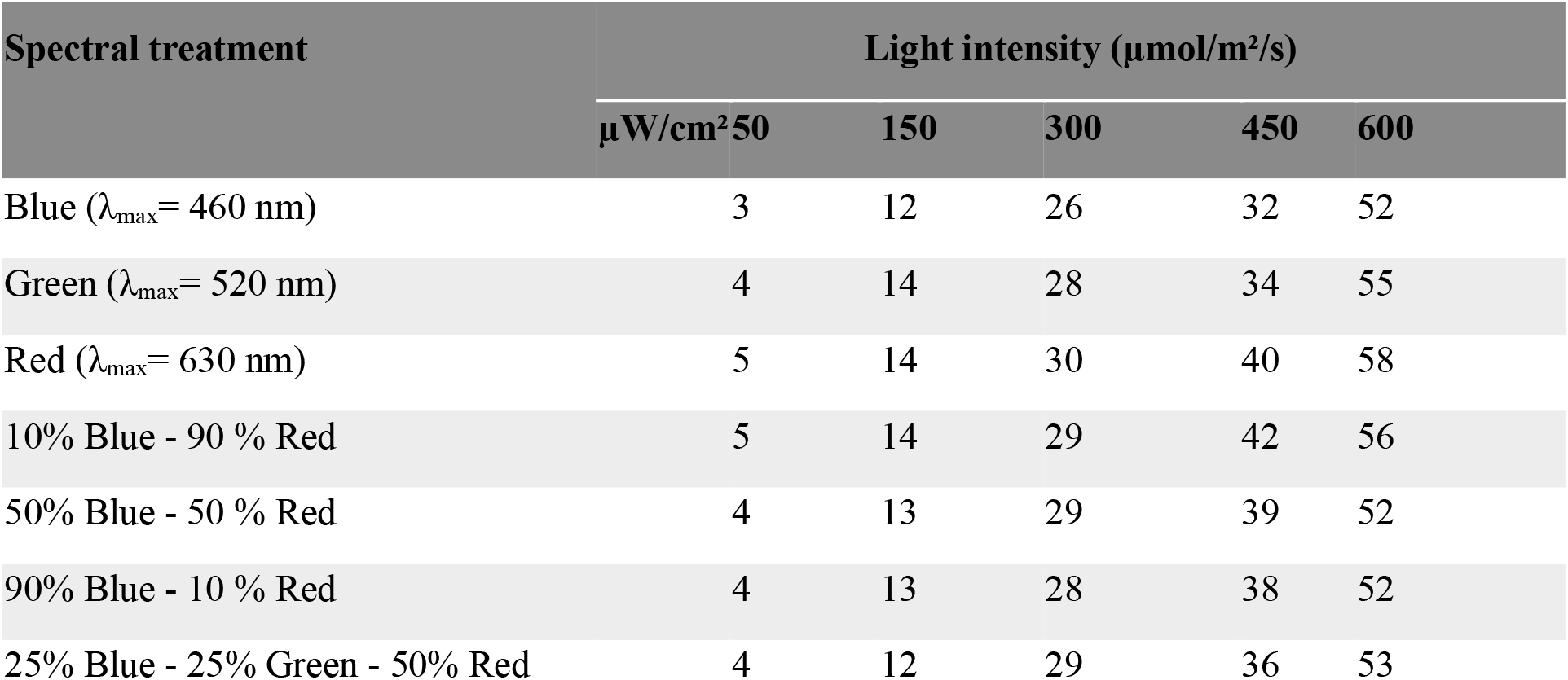
Light intensity values in PPFD (µmol/m^2^/s) for each lighting condition (spectral quality × intensity in µW/cm^2^).

To modulate the lighting environment for the microcosm experiments described in the next section, we developed a computer-controlled, spectrally agile optical device inspired by Cochard et al. (2017). This device was also optimized for high quality real-time video acquisition, to enable the analysis of predation behaviors at a small scale (1080p; 30 fps). Specifically, opaque chambers measuring 15 cm in height and 10.5 cm in diameter were equipped with a lid fitted with LEDs (product code B00K9M3WXG; frequency of 970 Hz) featuring a 2 cm opening for a camera attachment. The interior of the chamber was lined with aluminum. Three such chambers were connected and controlled in real-time from a laptop computer using a microcontroller (©ELEGOO UNO R3). A temperature probe (product code LYSB00Q9YBIJI-ELECTRNCS) was also integrated into each chamber, allowing for continuous data acquisition every 15 minutes. Each opaque chamber contained one experimental unit.

### Predation and locomotion behavior in microcosms

To assess the combined effects of various light spectra and intensities on *O. insidiosus* locomotion and predation behaviors against thrips on a small scale, we used female predators of the same age (synchronized emergence date ± 2–3 days), aged 3 to 10 days. They had continuous access to a source of water and food until they were used for the experiment, which took place in a dark room at ambient temperature. After being individually placed in transparent 1.5 mL microtubes (©Sarsted) their sex was determined under a stereomicroscope using morphological criteria (Isenhour & Yeargan 1981). A female *O. insidiosus* was then exposed to darkness for 30 minutes, followed by a 1-minute acclimation period to the tested spectrum × intensity combination in an opaque chamber. This procedure allowed us to isolate the effects of the tested artificial lighting treatment by minimizing the potential impact of rearing light conditions or ambient lighting (Cochard et al. 2017). The predator was then introduced into an experimental arena (experimental unit) containing 5 adult female thrips of similar size and color. The experimental arena consisted of clean and sterile Petri dishes 3.5 cm in diameter (©Fisherbrand™), covered with a Petri dish lid 5 cm in diameter (©Fisherbrand™) to prevent the thrips from escaping. In opaque chambers, only 5% of the light emitted by the optical device was reflected by the Petri dish lid, and the mean temperature was 25.5 ± 0.3 °C. The experimental units were randomized within each light treatment, which were also randomly assigned to a new opaque chamber at the beginning of each observation session. The behavior of the *O. insidiosus* and thrips was recorded for 5 minutes. Each *O. insidiosus* female was used for a single observation. Observations were conducted between 9:00 AM and 2:00 PM, with each of the 35 tested spectrum ×light intensity combinations tested twice per day for 7 consecutive days.

The experiment was repeated four times, involving four separate batches of predators to mitigate the potential influence of differences in vigor among shipments from the supplier, resulting in a final observation effort of 28 replicates per spectrum × intensity combination (140 replicates per spectrum).

The video recordings of these trials were analyzed using the open-source Behavioral Observation Research Interactive Software (BORIS) (Friard & Gamba, 2016). Locomotor activity, which influences predator-prey encounter probability, is significantly affected by the light spectrum and intensity in *O. sauteri* (Wang et al. 2013). However, previous studies have not examined interactions between *Orius* spp. and their prey, focusing instead on the optical manipulation for attraction (Park et al. 2023) or diapause avoidance (Herrick et al. 2021). Consequently, no data were available on attack probability or capture rates under artificial lighting, despite the importance of these behaviors for evaluating the control potential of *Orius* spp. in commercial settings. Therefore, the following behaviors were recorded: i) predator walking activity (time spent walking for more than 4 seconds); ii) attack attempts by the predator (the predator moved towards the prey, followed by direct contact between the predator and the targeted prey, either at the predator’s legs or rostrum); iii) successful capture (the predator immobilized and killed the targeted prey before inserting its rostrum to start feeding, corroborated by a dead thrips count at the end of observation).

### Predation in more complex arenas under 24-hour lighting sequences

To confirm the findings of the first experiment that predation of thrips by *O. insidiosus* occurs under a range of artificial lighting conditions, we tested a subset of the previously screened lighting treatments over a more realistic 24-hour schedule in a more complex experimental setup that also included plants. In the laboratory, three growth tents (©2022 VIVOSUN, Philadelphia, Ontario; L=121.9cm; W=121.9cm; H=182.9cm) were equipped with a dynamic spectrum luminaire (©2022 SOLLUM Technologies, Montreal, Quebec; model SF05-A), allowing real-time control of spectral quality and light intensity through the SUN as a Service® platform (“SUNaaS”). Ventilation (©2022 Inkbird, Shenzhen, China; model ITC-308 & ©2022 VIVOSUN, Philadelphia, Ontario, CA; model 43237-2) and humidification systems (©2022 Inkbird, Shenzhen, CHN; model IHC-200 & ©2022 ALACRIS, Ottawa, Ontario, CA; 4L humidifier) were used to maintain stable climatic conditions at 21 ± 2° C and 60 ± 2% RH. Based on the results of our microcosm experiment, we selected three artificial lighting spectra: 100% Blue (lowest proportion of prey captured), 25% Blue — 25% Green — 50% Red (intermediate proportion of prey captured), and 10% Blue – 90% Red (highest proportion of prey captured). To approximate conditions encountered in greenhouses at northern latitudes, we used these light treatments to extend the natural photoperiod in the late afternoon during light-limiting conditions—a technique known as ‘light supplementation’ (Runkle 2017) (Figure 1). Moreover, Ogino et al. (2020) found that the diurnal activity pattern of *O. sauteri* was maintained for 2 hours after sunset, and thus recommend this time of day to selectively attract this species to cropping systems using LEDs. In each treatment, “Sunlight” was artificially recreated natural solar spectrum, including a sunrise and sunset phase (λ_max_= 560 nm). These variations in light intensity and spectral composition over a 12-hour cycle were based on average spectrometric data provided by the World Meteorological Organization (WMO) for the Quebec City region and implemented in the SUNaaS platform (Honorine Lefevre, agr., SOLLUM Technologies inc., personal communication;). Since no effect of light intensity on small-scale predation performance of *O. insidiosus* was previously demonstrated between 3 and 58 µmol/m^2^/s, the trials were conducted at a constant intensity of 240 µmol/m^2^/s for the “Sunlight” period (mimicking a cloudy day) and about 30 µmol/m^2^/s for the different tested spectra.

**Figure 1.**
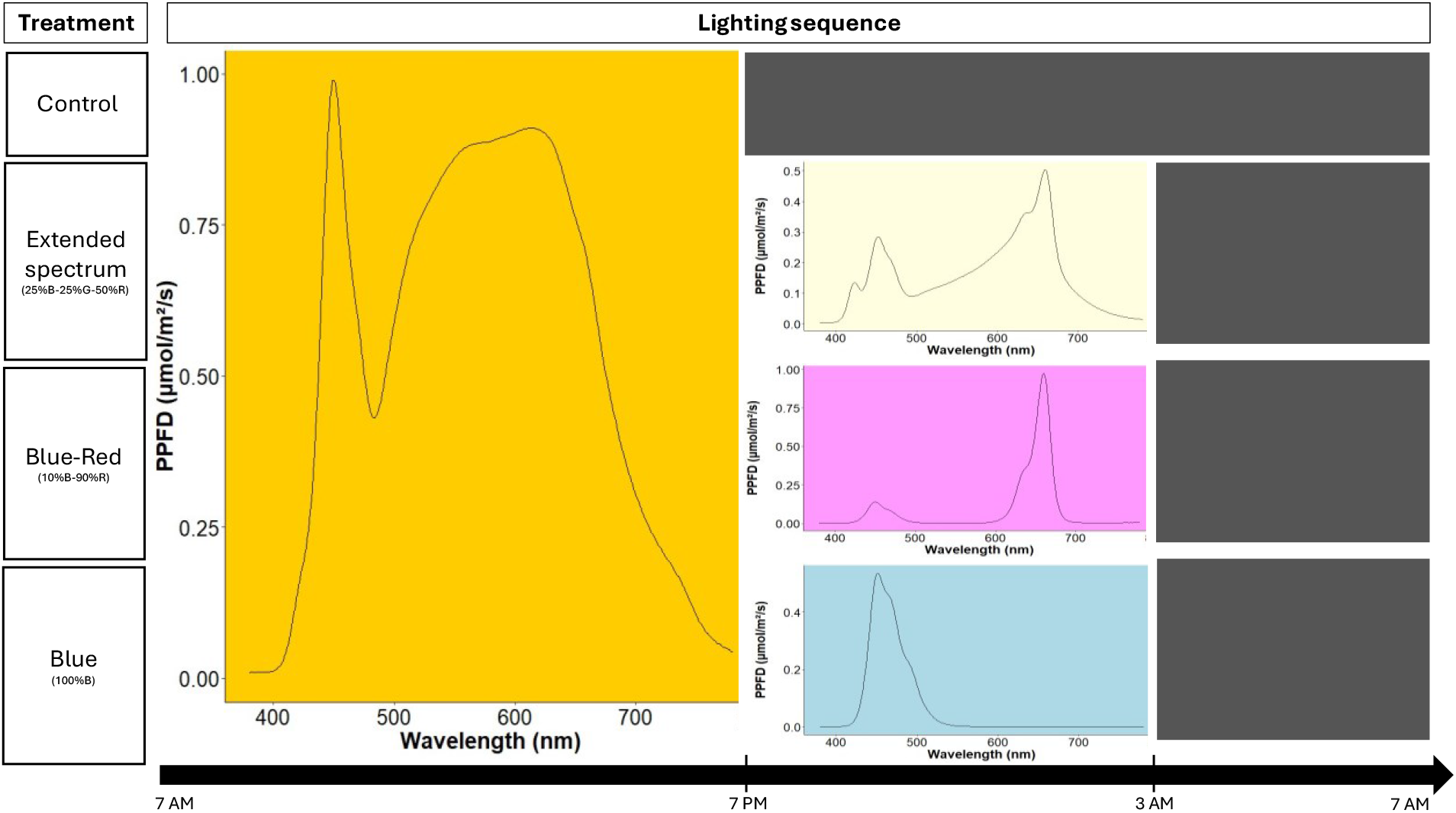
Lighting sequences tested in growth tents during the experiment in more complex arenas. The 24-hour cycle was divided into three parts: an initial phase of exposure to an artificially recreated ‘solar’ spectrum, followed by a second phase of photoperiod extension using one of the three tested light spectra (except for the control treatment), and finally, a dark phase (indicated by black rectangles). The light environment was modulated inside an opaque growth tent using a SOLLUM Technologies Inc. luminaire (model SF05-A), with a fixed light intensity of 240 µmol/m^2^/s for the ‘solar’ exposure phase (mimicking a cloudy day) and 30 µmol/m^2^/s for the photoperiod extension phase.

In protected crops, the artificial light spectrum can directly affect both plants and arthropods while also indirectly influencing arthropods through plant-mediated effects (Vänninen et al. 2010; Lazzarin et al. 2021). To account for this, prior to the experiment, the plants were exposed from the time of sowing to the same light sequence that would be tested during the trial, and the pest and predator insects were also acclimated to one of the tested sequences. The goal was to confirm the findings of the first experiment over the medium term in a more complex environment by assessing the capture performance of *O. insidiosus* against thrips, simulating a scenario where predators have acclimated to the light sequence for a few days, as would occur in protected crops following the introduction of natural enemies.

*Cucumis sativus* plants (Marketmore 70 variety, organic seeds; ©NORSECO, Laval, Québec, CA) were sown in 11.43 cm diameter pots containing a commercial growing mix (PRO-MIX BX; PremierTech Biotechnologies, Rivière-du-Loup, Québec, CA) and continuously exposed to one of the tested light sequences, with 6 plants per treatment. Progressive weekly fertilization (Plant-Prod 20-20-20 Classic; PlantProducts, Leamington, Ontario, CA) was initiated from the emergence of the true leaves.

Colonies of similarly aged thrips and *O. insidiosus* were placed in each of the tents (corresponding to a light treatment) for 10 days before they were used in experiments to allow acclimatization to the tested light sequence treatments. Colony care was conducted as described in the insects rearing section above. On the day of the experiment, cucumber plants aged 23– 25 days were transferred to 950 mL plastic transparent containers (©Elegant Disposables; experimental units) with modified lids to allow ventilation (8 cm diameter Nitex circle). Containers were then placed back into their original light treatment. Twenty minutes after the start of the phase corresponding to one of the three tested light spectra (100% Blue; 10% Blue-90% Red; 25% Blue-25% Green-50% Red), five adult female thrips were placed on each plant from the same light treatment and allowed to colonize it randomly for 20 minutes before introducing an adult female predator (age 10 ± 3 days), also from the same light treatment. The experimental units containing the plant, thrips, and a predator were then returned to their original light treatment for 2 hours before the predators were removed, and the number of live thrips was counted. The capture probability (whether at least one capture occurred or not) was then determined.

The experiment was repeated five times, involving five separate batches of predators, resulting in a final observation effort of 20 experimental units containing the tri-trophic system per tested light sequence. The experimental units were randomized within each light sequence, and the sequences were also randomly assigned to a new growth tent at the beginning of each temporal repetition.

For both experiments, the light environment was characterized using a LI-180 Spectrometer (LI-COR, Tucson, Arizona, USA), measured at Petri dish height in the experimental unit for the microcosm experiment and at plant height in the growth tent for the experiment in more complex arenas (Table 1; Figure 1).

### Statistical analysis

The data were analyzed using R software (R Core Team 2024; version 4.3.3), with a statistical significance threshold of p-values set at 0.05. Locomotor activity duration and the time before the first contact between the predator and one of its prey were subjected to square root and power transformations, respectively, as determined to be the most appropriate using the `boxcox` function from the `MASS` package (Venables & Ripley, 2002). These transformed data were then analyzed by linear mixed models *via* the `lme` function from the package `nlme` (Pinheiro et al., 2022). The probability of an attack and the proportion of successful captures (the number of captures divided by the number of attacks plus the number of captures) were analyzed using generalized linear mixed models with binomial error distributions, employing the `glmer` function from `lme4` package (Bates et al., 2015). For the experiment in more complex arenas, the number of captured prey was compared across spectra using the `glmer.nb` function from `lme4` package, while the capture probability was assessed with a generalized linear mixed model (binomial error distribution) using the `glmer` function. Multiple comparisons were conducted between the different levels of significant factors. In statistical analyses involving mixed models with random effects, the denominator degrees of freedom were sometimes approximated as infinite. This occurs because the tests rely on asymptotic distributions, where the degrees of freedom are not constrained by sample size and are treated as infinite to simplify calculations. Graphical representations illustrate the data adjusted from the full model, based on estimated marginal means computed with the `emmeans` function (Lenth, 2024), which derives least-squares means from the model, reflecting the influence of the covariates and factors considered.

All models were adjusted according to a generalized complete randomized block design, with temporal repetitions considered as blocks serving as repetitions of the different treatments, and each treatment also repeated within the same block. The conditions of normality and independence of residuals, as well as homoscedasticity, were verified. Analyses were validated by a professional statistician, Gaétan Daigle, MSc. (Statistical Consulting Service, Laval University).

## Results

### Predation and locomotion behavior in microcosms

The locomotion and predation behaviors of *Orius insidiosus* varied among some light spectra treatments, whereas light intensity tended to have negligible or marginal effects, their interaction being unsignificant in all cases (Table 2). The locomotor activity duration of adult female *O. insidiosus* varied with light spectrum but was not affected by light intensity (Table 2). Predators spent the most time moving under narrowband red, blue and green light conditions, while it was 1.6 times shorter when exposed to the 10% Blue - 90% Red treatment (Figure 2). The time before predators first contacted one of their prey varied with the light spectrum, but not light intensity (Table 2). Overall, the contact delay was less than 7 seconds across all spectra, with the shortest delay observed under narrowband green light. In comparison, it took 0.3 more time under red light and 50% Blue - 50% Red light ratio (Figure 2). The probability of *O. insidiosus* attempting to attack prey varied with light spectrum, with no effect of light intensity (Table 2). On average, the attack probability was greater than 70% and maximal values were associated with narrowband blue, green, and red lights, whereas this probability was 0.6 times lower for the 10% Blue - 90% Red ratio (Figure 2). Finally, the capture proportions (the number of captures divided by the number of attacks plus the number of captures) varied according to the light spectrum and light intensity (Table 2). We observed that a high number of attack attempts was not related to an increase in prey capture, as the narrowband spectra (which had the highest number of attack attempts) resulted in the lowest capture performance (Figure 3). The highest proportion of prey capture occurred under the 10% Blue - 90% Red treatment, while it was 1 and 1.75 times lower under narrowband red and blue, respectively. Meanwhile, the proportion of prey captured was reduced by a factor of only 0.2 when exposed to the extended spectrum relative to the 10% Blue - 90% Red treatment (Figure 3). Regarding light intensity, the highest proportion of prey was captured under 450 µW/cm^2^ across all spectra, while captures were reduced by a factor of 0.5 at 300 µW/cm^2^, although this difference was marginal (Figure 3).

**Table 2.** **Statistical comparison of locomotion and predator-prey behaviors of females *O. insidiosus* exposed to five prey in a Petri dish for 5 minutes, depending on light spectra, light intensity and their interaction (microcosm experiment).**

| Measurement | Factor | F-value<sub>df</sub> | P-value |
| --- | --- | --- | --- |
| Walking duration<sup>a</sup> | Spectrum | F<sub>(6, 932)</sub> = 16.90 | < 0.0001 |
|  | Intensity | F<sub>(4, 932)</sub> = 0.29 | 0.88 |
|  | Spectrum x intensity | F<sub>(24, 932)</sub> = 0.91 | 0.60 |
| Contact delay<sup>a</sup> | Spectrum | F<sub>(6, 642)</sub> = 2.71 | 0.01 |
|  | Intensity | F<sub>(4, 642)</sub> = 0.73 | 0.57 |
|  | Spectrum x intensity | F<sub>(24, 642)</sub> = 0.86 | 0.66 |
| Attack attempt probability<sup>b</sup> | Spectrum | F<sub>(6, ∞)</sub> = 9.02 | < 0.0001 |
|  | Intensity | F<sub>(4, ∞)</sub> = 0.55 | 0.70 |
|  | Spectrum x intensity | F<sub>(24, ∞)</sub> = 0.71 | 0.84 |
| Proportion of prey captured<sup>b</sup> | Spectrum | F<sub>(6, ∞)</sub> = 6.10 | < 0.0001 |
|  | Intensity | F<sub>(4, ∞)</sub> = 2.39 | 0.05 |
|  | Spectrum x intensity | F<sub>(24, ∞)</sub> = 1.42 | 0.09 |

a: analyzed using linear mixed model
b: analyzed using generalized linear mixed model with binomial error distribution

**Figure 2.**
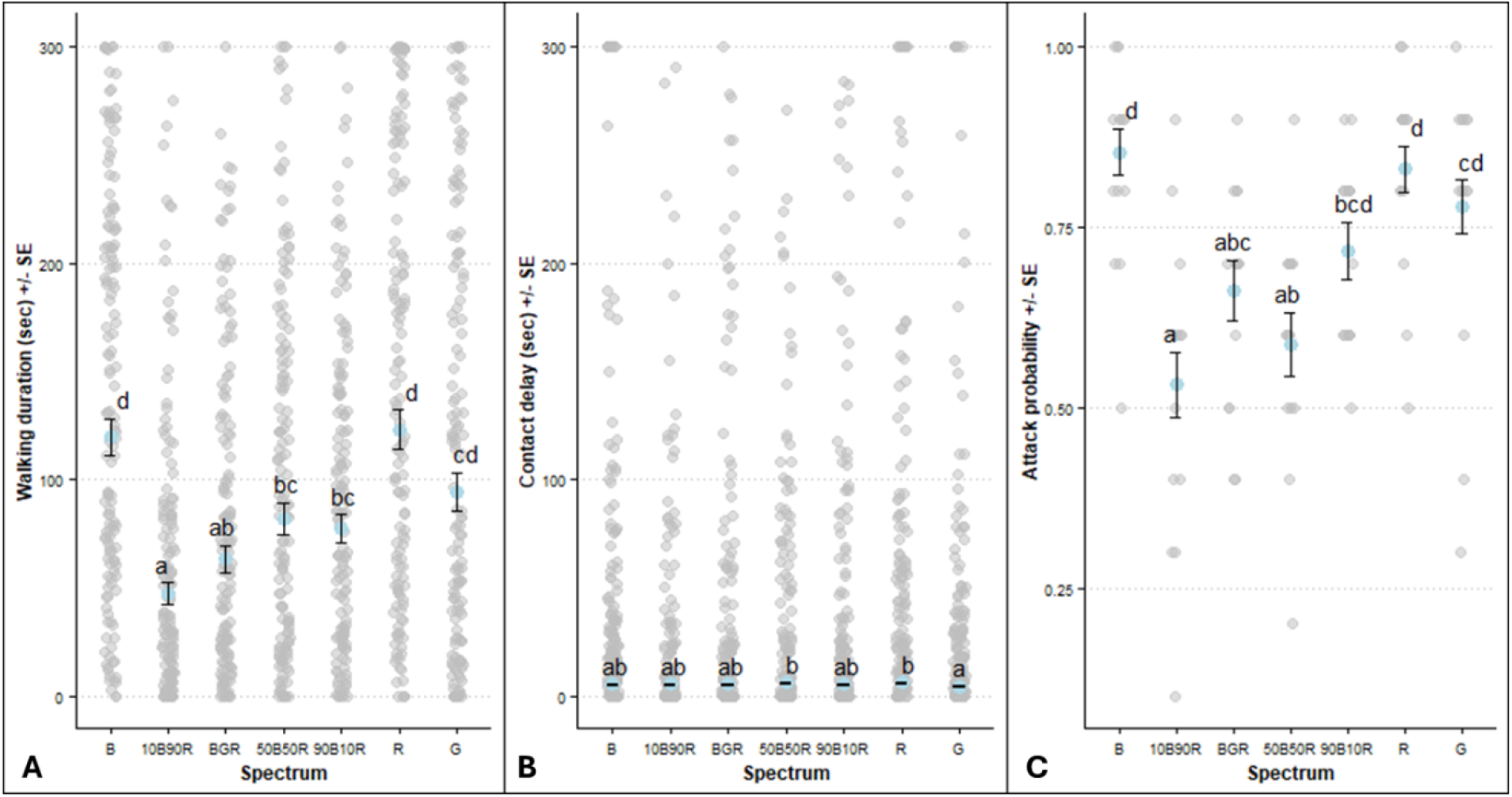
Behavioral responses of females *O. insidiosus* according to light spectra regarding A) Walking duration (± SE); B) Contact delay (± SE); C) Attack probability (± SE) during the microcosm experiment. The raw data are shown in gray, while the model-predicted means are in blue. The raw data for attack probabilities are derived from pooled observations for the same trial day.

**Figure 3.**
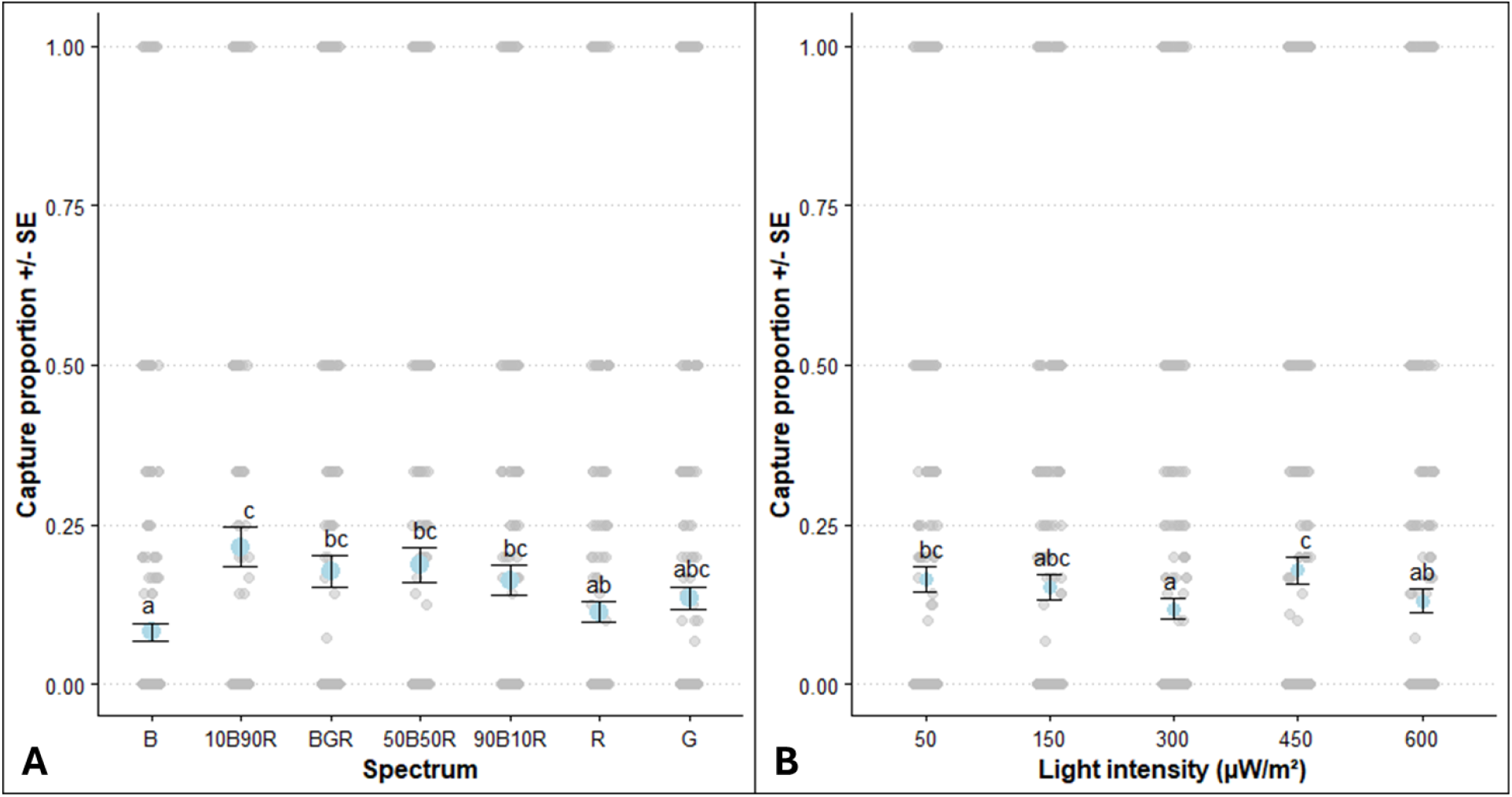
Capture proportions (the number of captures divided by the number of attacks plus the number of captures) (± SE) by adult female *Orius insidiosus* under different A) light spectra and B) light intensities during the microcosm experiment. The raw data are shown in gray, while the model-predicted means are in blue.

### Predation in more complex arenas under 24-hour lighting sequences

In growth tents, the number of surviving prey after 2 hours of exposure to an adult female *O. insidiosus* on cucumber plants partially corroborated observations made in our previous microcosm experiment, with captures being 0.38 times fewer under blue light (Figure 4). The same trend was observed for capture probability (i.e., whether at least one capture occurred or not), which varied with the light spectrum (Table 3). The highest capture probability was reached under the 10% Blue - 90% Red ratio, while it was reduced by a factor of 1.5 under narrowband blue light (Figure 4). However, increased capture rates did not translate to a significant difference in the number of surviving thrips among light spectrum treatments (Table 3).

**Table 3.** **Statistical comparison of capture performances of females *O. insidiosus* exposed to five prey on a cucumber plant for 2 hours, depending on light spectra (experiment in more complex arenas).**

| Measurement | Factor | F-value<sub>df</sub> | P-value |
| --- | --- | --- | --- |
| Number of surviving thrips<sup>a</sup> | Spectrum | F<sub>(2, ∞)</sub> = 2.34 | 0.10 |
| Capture probability<sup>b</sup> | Spectrum | F<sub>(2, ∞)</sub> = 4.45 | 0.011 |

a: analyzed using generalized linear mixed model with negative binomial error distribution
b: analyzed using generalized linear mixed model with binomial error distribution

**Figure 4.**
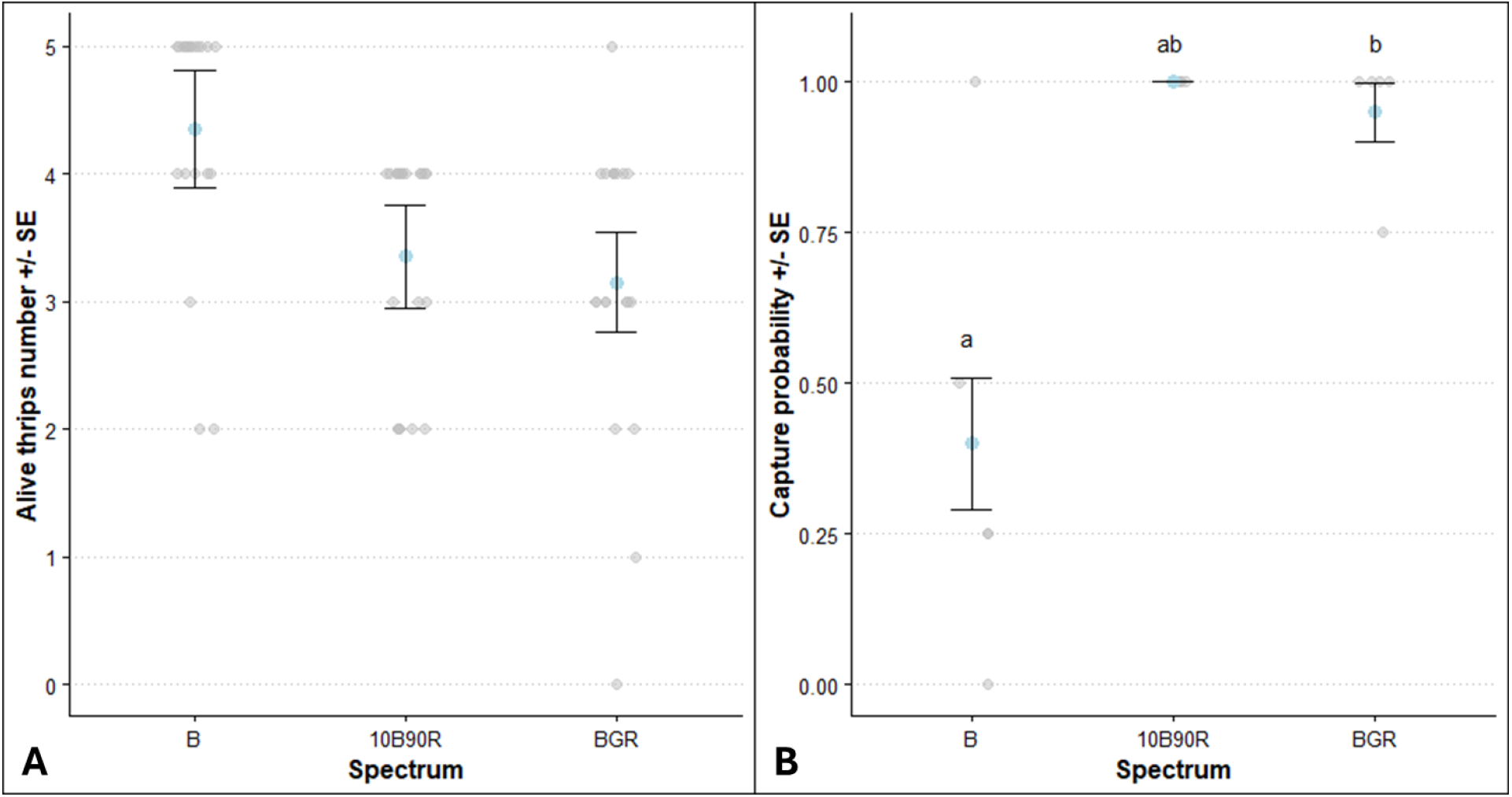
Number of surviving prey (± SE) (A) and capture probability (± SE) (B) after 2h-exposure to an adult female *Orius insidiosus* in arenas with plant under different light spectra during the experiment in more complex arenas. The raw data are shown in gray, while the model-predicted means are in blue. The raw data for capture probability are derived from pooled observations for the same temporal repetition.

## Discussion

*Orius insidiosus* successfully captured prey under all tested artificial lighting conditions, both in microcosms and complex arenas, even at low light intensities. The predator exhibited a high probability of attack across all spectra, quickly contacting prey in microcosms. However, the spectral composition, particularly the proportion of red light, significantly influenced predation behaviors. Notably, *O. insidiosus* was able to prey on thrips even in the absence of natural light in more complex arenas.

In the microcosm experiment, the light spectrum significantly influenced the predation behavior of *O. insidiosus*, while light intensity had minimal effect. Predation abilities remained consistent across intensities ranging from 3 to 58 µmol/m^2^/s. Unlike Wang et al. (2013), who found significant effects at 1000 to 3000 lux for *O. sauteri*, we detected no impact of light intensity on predator locomotion or capture rate, likely due to the narrower range and lower intensities we tested, constrained by our optical device. Our tested intensities were also lower than those used in commercial vegetable production (80 to 300 µmol/m^2^/s in Quebec). Bahşi and Tunç (2012) noted that *O. majusculus* can avoid diapause at light intensities as low as 10 lux, with minimal impact on its life cycle. Understanding how low light intensities support locomotion, predation, and development in *Orius* species can guide the selection of optimal lighting regimes for protected crops and the mass rearing of beneficials, ensuring that only the necessary amount of light is provided to natural enemies.

We observed that while *O. insidiosus* locomotion and predation varied under different monochromatic spectra, none fully impeded predation. Our predictions that blue and green lights would enhance predation, and red light would induce stress, were only partially validated. Contrary to expectations, short narrowband wavelengths (blue and green) resulted in low capture rates despite high attack probabilities. These wavelengths align with the visual sensitivity peaks of *F. occidentalis* (Matteson et al. 1992; Lopez-Reyes et al. 2022), potentially favoring prey escape. Given that thrips vision studies are mostly based on controlled trials, further research into their visual behavior and interactions with predators is needed (Lopez-Reyes et al. 2022). Red light caused high locomotor activity in *O. insidiosus* but low capture rates, suggesting an avoidance response, as predicted. Increased activity may have led to more encounters, but insufficient energy allocation during attacks may have hindered the predator’s success, possibly due to thrips’ defensive behaviors (Cox et al. 2006; M. L. Canovas personal observation). Similarly, Wang et al. (2013) reported that red light increased walking speed and respiration in *O. sauteri*, linked to an avoidance response, and negatively affected pre-oviposition delay, fecundity, and development. Interestingly, UV light hindered *O. insidiosus* from capturing prey, contradicting the positive phototaxis seen in *O. sauteri* (Ogino et al. 2015; 2016; 2020; Park et al. 2023) and *Nesidiocoris tenuis* (Uehara et al. 2019; Park and Lee 2021; Park et al. 2022; 2023). Our results suggest that visual ecology studies of beneficials should consider prey interactions to assess the net impact of light on biological control. The observed interspecific variability under UV wavelengths also underscores the need to further explore visual ecology in commercially important beneficial species.

We observed varied responses of *O. insidiosus* to mixed spectra, challenging our initial predictions that blue light would be beneficial and red light would induce stress. Surprisingly, the 10% Blue - 90% Red spectrum significantly increased capture success compared to narrowband red, despite lower attack probability and reduced locomotor activity. This ratio appears optimal for biological control, potentially balancing the negative effects of narrowband blue and red by creating a favorable visual environment for *O. insidiosus* predation on *F. occidentalis*. We hypothesize that this spectrum might be less detectable by the prey’s visual system or mitigate red light-induced stress in the predator, as suggested above. Low locomotor activity likely reflects the predator remaining stationary while consuming prey (M. L. Canovas, personal observation across spectra). The behavior of abandoning carcasses for live prey, as reported by Cox et al. (2006), was not observed during our 5-minute trials. The extended spectrum (25% Blue - 25% Green - 50% Red) showed attack probabilities and capture proportions similar to or slightly lower than those of the 10% Blue - 90% Red treatment. This result is promising given the common use of various Blue–Green–Red light ratios in horticulture (Orlando et al. 2022), where green wavelengths support photosynthesis (Smith et al. 2017) and improve human detection of phytopathological symptoms. Increasingly, studies on beneficials’ behavior under artificial lighting are testing extended Blue–Green–Red spectra beyond traditional blue-red ratios (Fraser et al. 2023a; 2023b; Gonzalez et al. 2023). Our tested extended spectrum closely matches that identified by Garcia and Lopez (2020) as enhancing transplant vigor and biomass in greenhouse vegetable crops. Overall, our findings on mixed lighting ratios suggest that it is realistic to adjust lighting regimes to balance agronomic productivity with biological control, optimizing electrical resource use while maintaining the effectiveness of natural enemies and supporting sustainable agriculture in the context of climate change.

*O. insidiosus* successfully captured thrips under sole-source artificial lighting in complex arenas with plants, regardless of the light sequences tested. The reduced capture proportion under blue light observed in microcosms was also reflected by a lower capture probability in complex arenas compared to mixed spectra. However, prey survival rates remained similar across all lighting treatments, possibly due to the short exposure duration. As *O. insidiosus* is diurnal and kills around 22 thrips in 24 hours (Tommasini et al. 2004; Silva et al. 2022), the observed rates in mixed spectra align with expected values. A longer exposure might reveal a significant reduction in prey consumption under blue light. Predator satiation, given continuous food access before the experiment, and the filtering of artificial light by the plant canopy may have further reduced the impact of lighting on predation behaviors compared to microcosms. This study is the first to examine interactions between *O. insidiosus* and its prey under artificial lighting in both microcosms and complex arenas. Building on established practices, laboratory screening should identify promising light treatments for natural enemies (Ogino et al. 2015; Cochard et al. 2017; Fraser et al. 2023a), which can then be tested in more commercially relevant conditions involving plants (Ogino et al. 2016; Cochard et al. 2019b; Fraser et al. 2023b; Canovas et al., in preparation).

Our results support previous findings that beneficial insects can effectively control pests under various light environments, including solely artificial sources. This knowledge is crucial not only for optimizing the mass rearing of beneficials but also for adjusting horticultural light spectra in greenhouses and indoor farming. Transitioning biological control from greenhouses to fully indoor cultivation, with its enclosed spaces and sole-source artificial lighting, introduces new challenges (Roberts et al. 2020). Notably, lighting management is one of the key factors influencing biological control efficiency in commercial indoor production (Lemay et al. 2022). The impact of artificial lighting on tri-trophic interactions—among predators, prey, and plants—has significant implications for biological control outcomes (Väninnen et al. 2011; Lazzarin et al. 2021; Meijer et al. 2023a; 2023b). Therefore, larger-scale lighting experiments in more realistic horticultural systems are necessary.

## Acknowledgments

The authors would like to acknowledge the partnership and financial support received from the Ministère de l’Agriculture, des Pêcheries et de l’Alimentation du Québec, MITACS, SOLLUM Technologies inc., ANATIS Bioprotection and the National Optics Institute (project n° IA120649 & IT17200). The authors would like to specifically thanks Patrick Larochelle (Tech. COPL), Jean-Michel Vigneault (BSc. student), Gaétan Daigle (MSc. Laval University) and Sidy Bouya Ndiaye (M. Ing) as well as Jonathan Bélanger (Tech.) (SOLLUM Technologies inc.) for their technical support.

## Authors’ contribution

MLC, TG, JFC and MD secured the funding; MLC, TG, and JFC designed the optical device; MLC designed and conducted the experiment, then analyzed and interpreted the data; MLC and PKA wrote the first draft of the manuscript; MLC, PKA, TG, JFC, and MD revised and edited the text.

## Statements and Declarations

### Ethical approval

The authors approve the ethical guidelines of the journal.

### Consent for publication

All authors gave consent to the publication of the study.

---

**Supplementary Material 1.**
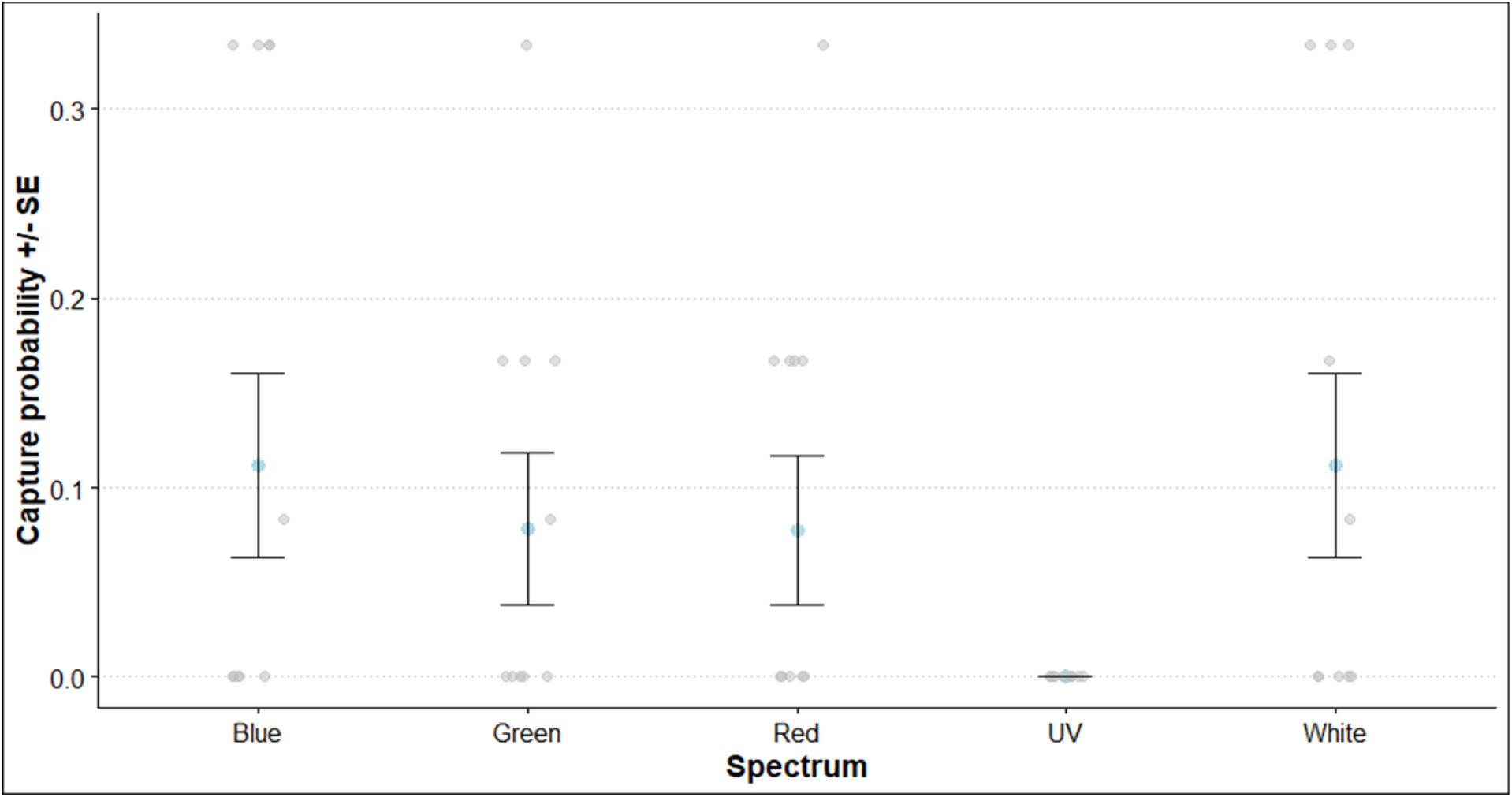
Capture probability (± SE) by adults *Orius insidiosus* during preliminary trials. The capture probability (± SE) by adult *O. insidiosus* during preliminary trials did not significantly vary according to light spectrum (F _(4, ∞)_ = 0.193, p = 0.9421), predator’s sex (F _(4, ∞)_ = 0, p = 1), or their interaction (F _(4, ∞)_ = 0.145, p = 0.9653). Interestingly, no predation was observed for the UV treatment (λ_max_ = 400 nm; product LED code: B0B9BMBV1N). Using the same methodology as described in Predation and locomotion behavior in microcosms section, behaviors of adult female and male *O. insidiosus* from the same batch were recorded during 5-minute independent Petri dish observations of one predator and five prey (n = 79 observations per spectrum). Light intensities tested were 10, 160, and 320 µW/m^2^, and the White spectrum was composed of 33% Blue, 33% Green, and 33% Red lights. Capture probability was analyzed using mixed binomial models with the `glmmTMB` function from `glmmTMB` package (Brooks et al., 2017), specifying a binomial distribution (α = 0.05). The raw data are shown in gray, while the model-predicted means are in blue. The raw data are derived from pooled observations for the same temporal repetition.

